# Cellular source and circuit context organize functional specificity in the *Drosophila* NPF system

**DOI:** 10.64898/2026.09.08.750118

**Authors:** Madison N. Endres, Tarandeep Singh Dadyala, Kevin W. Christie, Irina T. Sinakevitch, Lisha Shao

## Abstract

Neuropeptides regulate diverse and sometimes opposing functions, yet how a single peptide achieves functional specificity remains unclear. Receptor diversity can provide specificity, but not in systems with only one receptor. *Drosophila* neuropeptide F (NPF) acts through a single receptor, providing a model to test whether specificity is organized by the peptide’s cellular source. Using the whole-brain connectome, we defined four adult NPF cell types with no shared first-order synaptic partners and separate networks. These cell types made distinct and partly opposing contributions to reinforcement, feeding, mating, and energy storage. Most peptide-dependent effects were undetectable after population-level NPF reduction, showing that global perturbation can obscure source-specific functions. Changing neuronal activity and reducing NPF in the same cells produced different, sometimes opposing, phenotypes, so neuronal activation cannot be equated with peptide action. Thus, cellular source and circuit context generate functional specificity that population-level perturbation conceals.

## Introduction

Neuropeptides act as neuromodulators that reconfigure neural circuits according to an animal’s internal state and social context ^1^. Yet individual neuropeptides often regulate a remarkably broad range of physiological and behavioral processes. How a shared peptide signal achieves functional specificity across these diverse outputs remains a central problem in neuropeptide biology ^2,3^. A paradigmatic example is Neuropeptide Y (NPY), an evolutionarily conserved brain-gut peptide with established roles in feeding, energy metabolism, sleep, stress, reward processing, and reproductive physiology ^4–7^. This breadth poses an immediate conundrum: how can a single peptide regulate so many, and sometimes opposing, functions with specificity?

One explanation is receptor diversity: NPY acts through multiple receptor subtypes with distinct expression patterns and signaling properties ^8^, allowing the same ligand to act differently on different targets. This mechanism cannot explain functional diversification in peptide systems that signal through a single receptor, such as nociceptin/orphanin FQ.

Another complementary mechanism is cellular-source specificity: peptide-expressing cell types may occupy distinct circuit positions, allowing the same peptide to influence different downstream networks ^2,3^. Yet it remains unclear how peptide-dependent functions are partitioned across these cell types, whether their contributions are concordant or opposing, and how they are represented by population-level perturbation.

Addressing this question in mammals is challenging because neuropeptide-expressing neurons are typically broadly distributed and embedded in complex circuits. In contrast, the genetic and anatomical tractability of the fruit fly *Drosophila melanogaster* offers an opportunity to resolve these questions at cell-type resolution. *Drosophila* neuropeptide F (NPF) is a homolog of the vertebrate NPY family ^9^ but, unlike NPY, signals through a single known receptor, NPFR ^10^. Receptor-subtype identity therefore cannot account for the diversification of NPF function, making NPF a tractable system in which to test whether cellular source and circuit context contribute to functional specificity.

Despite this simpler receptor architecture, NPF regulates as broad a range of functions as NPY. It is classically orexigenic: *npf* expression is associated with food attraction in larvae ^11^, and NPF signaling enables food-deprived animals to overcome aversion and consume noxious food ^12^. Consistent with this role, satiety-associated signals suppress feeding drive by inhibiting NPF neurons ^13,14^. Beyond feeding, NPF gates hunger-dependent expression of appetitive memory through mushroom-body dopaminergic neurons ^15^, promotes starvation-induced wakefulness ^16^, and links sexual experience to reward state, metabolic physiology, stress resistance, and lifespan^17^.

These findings also expose an apparent tension in how NPF has been interpreted across reward contexts. In feeding and memory, increased NPF signaling is associated with deprivation-driven food seeking and memory expression, consistent with a state in which reward is lacking ^11,12,15^. In male sexual experience, by contrast, successful copulation increases brain NPF, whereas the reduced NPF-system activity that results from sexual deprivation increases ethanol seeking ^18^. Intriguingly, activation of NPF neurons is itself rewarding and can reduce subsequent ethanol seeking ^18,19^, consistent with a state of reward attainment ^18^. These observations arise from different sexes, developmental stages, cell populations, behavioral contexts, and modes of perturbation; apparently opposing phenotypes are therefore not directly contradictory. Nevertheless, they are difficult to accommodate within a unified interpretation of NPF signaling. This raises a specific question: are hunger- and mating-related functions carried by distinct NPF cell types with different circuit outputs? Moreover, it remains unknown how adult NPF cell types are organized across the connectome, how their peptide-dependent functions compare across behaviors and physiology, and whether population-level manipulation preserves or obscures their individual contributions.

In this study, we asked how functional specificity is organized across distinct adult NPF cell types within a one-peptide-one-receptor system and how their individual contributions are represented when the peptide system is manipulated at the population level. We constructed a functional map of the NPF population by combining the *Drosophila* whole-brain connectome with cell-type-specific genetic manipulation and quantitative behavioral and physiological assays. We defined four sexually monomorphic NPF cell types by morphology and connectivity, generated or obtained drivers that provided genetic access to each population, and compared the effects of cell-type-specific NPF knockdown with neuronal activation or silencing across mating, feeding, reward, learning, and metabolic assays.

We found that distinct NPF cell types occupied largely separate connectomic positions and produced separable phenotypes. Population-level NPF loss failed to reveal most cell-type-specific peptide-dependent phenotypes, whereas neuronal activity manipulations frequently produced effects distinct from NPF knockdown. These findings identify cellular source, circuit connectivity, and signaling repertoire as complementary organizers of functional specificity in the absence of known receptor-subtype diversity.

## Results

Anatomically distinct NPF subsets have been linked to reward ^19^, feeding motivation ^20^, larval sugar-reward learning ^21^, and satiety ^13^. A recent anatomical survey further resolved approximately 50 adult NPF neurons into five morphological clusters, including ventrolateral neurons that project centrifugally to the optic lobes ^22^. Yet despite this progress, a systematic, connectome-guided, cell-type-resolved functional map of the adult NPF population was lacking. To resolve this gap, we combined the whole-brain connectome (FAFB/FlyWire) ^23,24^ with NeuronBridge ^25^ and available split-GAL4 resources ^26^ to define NPF cell types by morphology and connectivity, establish cell-type-specific genetic access, and compare their functions.

### NPF cell types possess distinct connectomic networks

To characterize the organization of the adult NPF population, we identified four sexually monomorphic NPF-expressing cell types (hereafter NPF cell types) in the FAFB whole-brain connectome ^23,24^ (Table S1). We named them dorsomedial (DM), lateral type 1-large (L1-l, referred to as L1 hereafter), posterior type 1 (P1; unrelated to the *fruitless⁺* P1 courtship-command neurons), and posterior type 2 (P2). The four cell types differed in morphology and projection pattern (Figure 1). We detected no synapses between different NPF cell types, although the bilateral P2 neurons formed within-type connections (Figure S1A).

**Figure 1.**
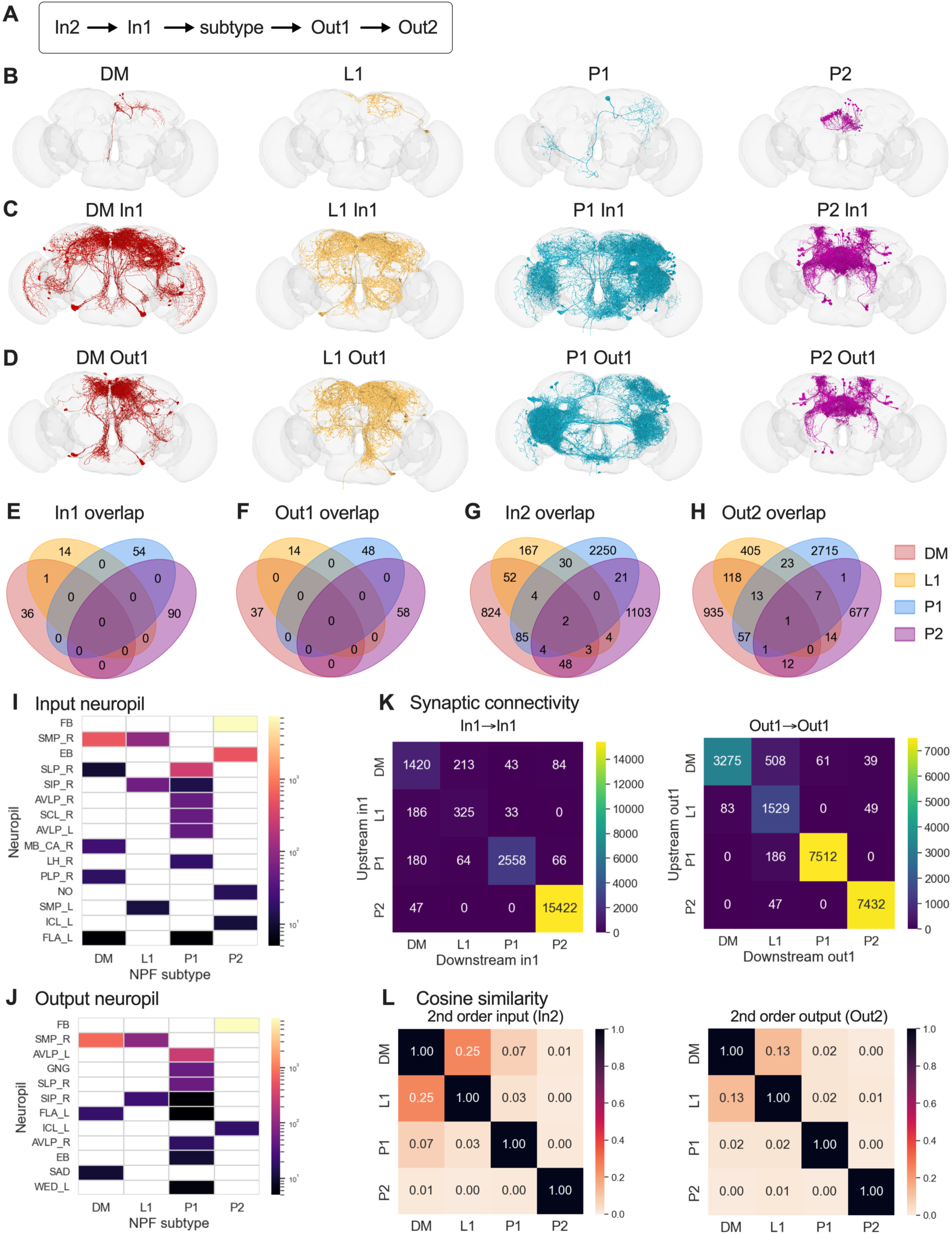
NPF cell types have distinct connectivity in the connectome. (A) Schematic defining the orders of synaptic partners relative to each NPF subtype; arrows indicate the direction of information flow. In1, first-order synaptic input (direct presynaptic partners); In2, second-order synaptic input; Out1, first-order synaptic output (direct postsynaptic partners); Out2, second-order synaptic output. Connectome-based annotations predict that DM and P1 are cholinergic, whereas L1 and P2 are serotonergic. (B) Morphology of the four monomorphic NPF cell types—DM, L1, P1, and P2. Only cells in the right hemisphere are shown for clarity. Subtypes are color-coded throughout the figure (DM, red; L1, yellow, P1, teal; P2, magenta). (C) Morphology of the first-order presynaptic (input) partners (In1) of each NPF cell types, shown for the right hemisphere. (D) Morphology of the first-order postsynaptic (output) partners (Out1) of each NPF cell types, shown for the right hemisphere. (E-H) Venn diagrams showing the overlap in synaptic partners among the four NPF subtypes for (E) first-order inputs (In1), (F) first-order output (Out1), (G) second-order inputs (In2), and (H) second-order outputs (Out2). Numbers indicate the count of unique or shared partner neurons. (I and J) Heatmaps showing the neuropil distributions of inputs (I) and outputs (J) for each NPF subtype. FB: fan-shaped body; SMP: superior medial protocerebrum; EB: ellipsoid body; SLP: superior lateral protocerebrum; AVLP: anterior ventrolateral protocerebrum; SCL: superior clamp; MB_CA: mushroom body calyx; LH: lateral horn; PLP: posterior lateral protocerebrum; NO: noduli; SIP: superior intermediate protocerebrum; ICL: inferior clamp; FLA: flange. (K) Connectivity between first-order input partner sects (In1◊In1, left) and between first-order output partner sects (Out1◊Out1, right). Color indicates total synapse number. (L) Pairwise cosine similarity between subtypes based on synapse-weighted second-order partner vectors for second-order inputs (In2, left) and outputs (Out2, right). 0, no shared partners; 1, identical partners; diagonal values are 1 by definition.

We next mapped the first- and second-order presynaptic and postsynaptic partners of each cell type (Figure 1A, C, D). The four cell types had largely non-overlapping first-order presynaptic partners, with only one partner shared between DM and L1 (Figure 1E), and completely non-overlapping first-order postsynaptic partners (Figures 1F and S1C), whereas their second-order partner sets overlapped only modestly (Figure 1G, H). This segregation was also evident at the neuropil level. The only input neuropils shared by multiple cell types were superior medial protocerebrum (SMP; DM and L1), superior lateral protocerebrum (SLP; DM and P1), superior intermediate protocerebrum (SIP; L1 and P1), and weakly flange (FLA; DM and P1) (Figure 1I), whereas the only shared output neuropils were the SMP (DM and L1) and SIP (L1 and P1) (Figure 1J).

Distinct partner identities do not preclude communication between their broader networks. We therefore asked whether the first-order partners of one NPF cell type also contacted other NPF cell types and whether the partner populations of different NPF cell types synapsed onto one another. First-order partners contacted their corresponding NPF cell type but not the other NPF cell types (Figures S1B and S1C), and synapses between partner populations were sparse and pair-specific (Figure 1K). Pairwise cosine similarity, which incorporates synaptic weight rather than partner membership alone, likewise showed negligible similarity among first-order partner vectors (Figure S1D) and low similarity among second-order vectors, with the highest values between DM and L1 (Figure 1L).

Together, these analyses show that DM, L1, P1, and P2 are embedded in largely distinct input-output networks. DM and L1 were the most similar pair at the second-order level, but their direct partners and overall connectivity remained separable.

### Distinct NPF cell types differentially recruit PAM dopaminergic neurons

Anatomical segregation suggests, but does not establish, functional specialization. In previous work using stochastic genetic intersection, we found that optogenetic activation of DM neurons, but not the other NPF populations tested, induced positional preference ^19^.

We therefore sought stable genetic access to each cell type for systematic functional comparison. Using the FAFB connectome, NeuronBridge, and GAL4 and split-GAL4 resources ^23–26^, we generated stable drivers for DM, P1, and P2 and obtained an existing driver for L1 ^27^. Immunostaining confirmed that each driver labeled the intended NPF cell type in the central nervous system (Figures 2A and S2A). The drivers showed little or no gut expression, except for weak labeling by the P2 driver in a restricted gut region (Figure S2B). These drivers allowed us to compare the consequences of reducing NPF or manipulating neural activity in each cell type.

**Figure 2.**
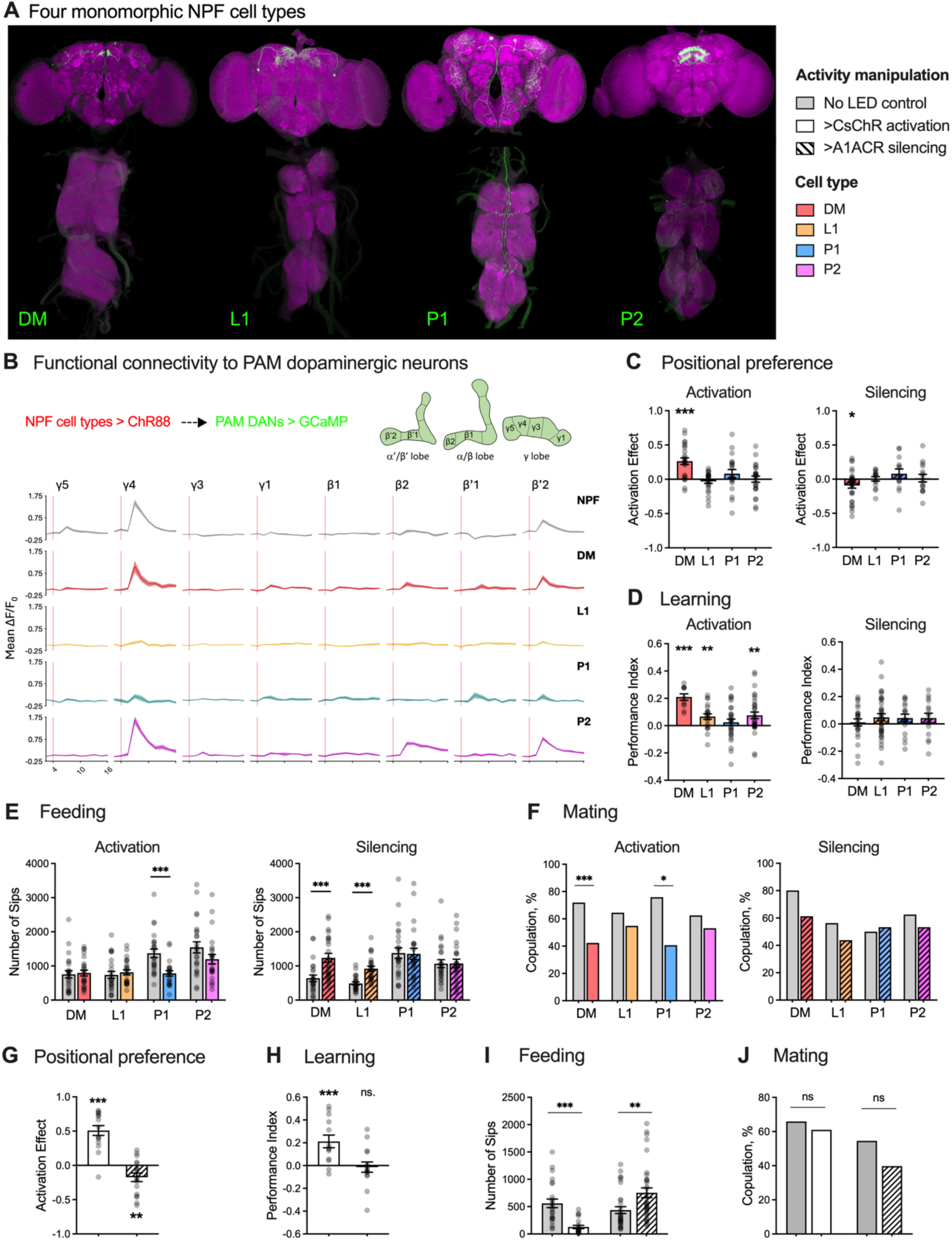
Cell-type-specific and population-level activity manipulations reveal functional specialization within the NPF system. (A) Expression patterns of the split-GAL4 (SS) driver for each NPF subtype in the brain (top) and ventral nerve cord (bottom), revealed by immunostaining of SS-driver>CsChrimson flies. Green, anti-GFP; magenta, anti-nc82 (neuropil); scale bar, 100 µm. (B) Functional connectivity to PAM dopaminergic neurons (DANs), measured by recording calcium signals from PAM DANs while optogenetically stimulating each NPF subtype (SS-driver>Chrimson88; 58E02>GCaMP7s). Traces show mean ΔF/F per PAM compartment. Activating DM or P2 neurons evoked calcium responses in PAM DANs innervating the γ4 and β′2 compartments, recapitulating the pattern evoked by the full NPF-GAL4 driver. NPF subtypes were optogenetically activated (SS-driver>CsChrimson) or silenced (SS-driver>A1ACR). In panels C-F, the left subpanel shows activation and the right subpanel shows silencing; on-LED siblings served as controls. (C) Activating DM neurons produced positional preference for optogenetic stimulation zone, whereas silencing DM produced avoidance. (D) Pairing an olfactory conditioned stimulus with activation of DM, L1, or P2 generated appetitive memory (positive performance index), whereas silencing did not generate memory. (E) Activating P1 neurons decreased food consumption, whereas silencing DM or L1 neurons increased it. (F) Activating DM and P1 neurons reduced female copulation percentage, whereas silencing the individual subtypes did not affect copulation percentage. (G-J) All NPF-expressing cells were globally activated (NPF-GAL4>CsChrimson) or silenced (NPF-GAL4>A1ACR), with no-LED siblings as controls. (G) Global activation produced positional preference for the optogenetic stimulation zone, whereas global silencing produced avoidance. (H) Pairing an olfactory conditioned stimulus with global activation generated appetitive memory; global silencing did not. (I) Global activation suppressed food intake, whereas global silencing increased it. (J) Neither global activation nor global silencing affected female copulation rate. N = 15-25 (positional preference), 11-35 (learning), 22-24 (feeding), n = 29-59 (mating). Data are mean ± SEM. Statistics: Fisher’s exact test (mating); unpaired *t*-test or Mann-Whitney test (feeding); one-sample *t*-test or Wilcoxon signed-rank test (positional preference and learning), the criterion for choosing parametric vs non-parametric tests depending on normality by Shapiro-Wilk. \**p* < 0.05, \*\**p* < 0.01, \*\*\**p* < 0.001.

We first asked whether the four NPF cell types differentially recruit a canonical appetitive reinforcement pathway, the function most consistently attributed to NPF. In adult flies, activation of NPF-expressing neurons can reinforce an associated odor ^18,19^. NPF signaling also regulates the expression of food-reward memory through mushroom-body-innervating dopaminergic neurons ^15^, raising the possibility that defined NPF cell types recruit dopaminergic reinforcement pathways. Because protocerebral anterior medial (PAM) dopaminergic neurons mediate positive reinforcement and drive compartment-specific synaptic plasticity in the mushroom body during appetitive learning ^28^, we tested whether activating individual NPF cell types evokes responses in PAM axons.

We optogenetically stimulated each NPF cell type while recording calcium signals from PAM axons that innervate specific compartments of the mushroom body horizontal lobes. Activating DM or P2 neurons evoked calcium responses in PAM axons that innervate the γ4 and β’2 compartments, recapitulating the compartment pattern observed after pan-NPF activation (Figure 2B). In contrast, activating P1 or L1 did not evoke a detectable calcium response in the PAM compartments examined.

These results place DM and P2 neurons functionally upstream of the PAM dopaminergic neurons examined and show that NPF cell types have distinct access to PAM reinforcement circuitry. Because the connectome contained no direct synapses from the four NPF cell types onto PAM neurons, it remains unclear whether DM and P2 recruit PAM through polysynaptic pathways or by peptidergic signaling.

### Neural activity manipulations reveal cell-type-specific behavioral functions

We next tested whether differential access to reinforcement circuitry was accompanied by distinct behavioral effects. We first asked whether acutely activating or silencing each NPF cell type carried positive or negative valence in a positional preference assay ^19^. Flies were allowed to choose between an unstimulated zone and a zone in which a selected cell type was optogenetically activated with CsChrimson ^29^ or silenced with A1ACR ^30^. In this assay, preference for the stimulation zone indicates that the manipulation is appetitive, whereas avoidance indicates that it is aversive.

Activating DM induced positional preference for the stimulation zone, whereas silencing DM induced avoidance. In contrast, manipulating L1, P1, or P2 had no detectable effect on positional preference (Figure 2C). Thus, among the four cell types tested, only DM manipulation altered positional valence.

We then asked whether the valence generated by manipulating neural activity could reinforce an associated olfactory cue. In olfactory conditioning, pairing an odor with activation of DM, L1, or P2 neurons generated appetitive memory (Figure 2D), indicating that each population is sufficient to provide, or recruit, a positive reinforcing signal. By contrast, silencing any individual NPF cell type during odor presentation generated neither appetitive nor aversive memory under these conditions (Figure 2D). Notably, L1 activation supported appetitive memory despite producing no detectable positional preference and no detectable response in the PAM compartments examined. Thus, immediate preference and appetitive reinforcement are supported by overlapping but nonidentical NPF cell types.

The preceding assays used artificial neural stimulation as the valenced or reinforcing event. Because NPF signaling also regulates behaviors directed toward natural rewards, such as food and mating opportunities ^11,16,31,32^, we next asked whether the four cell types differentially influence food intake and copulation.

We quantified food consumption during cell-type-specific activation or silencing using optoPad ^33–35^. Activating P1 decreased consumption, whereas silencing DM or L1 neurons increased it (Figure 2E). To assay mating, we paired wild-type males with individual experimental females during optogenetic stimulation. Activating DM or P1 neurons reduced female copulation rate, whereas silencing any individual NPF cell type had no detectable effect (Figure 2F). These results extend the functional segregation of NPF cell types from artificial reinforcement to natural reward-related behaviors.

Having established a cell-type-resolved activity map, we next asked how these contributions combine when the entire NPF population is manipulated. If population-level activity simply aggregated the effects of individual cell types, the direction of cell type-specific phenotypes should be retained; departures from that pattern could instead reflect interactions or nonlinear integration across the population.

Globally activating NPF-expressing cells induced positional preference, whereas global silencing led to avoidance (Figure 2G), matching the directions of DM activation and silencing (Figure 2C). Global activation generated appetitive olfactory memory, whereas global silencing did not support memory formation (Figure 2H), consistent with the effect of activating DM, L1, or P2 individually (Figure 2D). Global activation decreased food intake, whereas global silencing increased it (Figure 2I), matching the directions of P1 activation and DM or L1 silencing, respectively (Figure 2E). In contrast, neither global activation nor silencing affected copulation rate (Figure 2J), despite the reduction produced by activating DM or P1 individually (Figure 2F).

Thus, manipulating activity at the population level preserved the direction of the cell-type-specific effect in positional preference, learning, and feeding, but not mating. The mating exception shows that population-level manipulation does not invariably reveal the effects of its constituent cell types and may instead reflect differential recruitment or nonlinear interactions among populations.

Importantly, however, these activity manipulations do not identify the contribution of NPF itself: optogenetic activation or silencing changes the neurons’ aggregate activity-dependent output, including NPF and any co-released fast-acting transmitters. To distinguish NPF-dependent effects from those of the broader neuronal output, we next reduced NPF selectively within each cell type using cell-type-restricted RNAi against *npf*.

### Cell-type-specific NPF knockdown reveals distinct peptide-dependent functions

The positional-preference and conditioning assays above rely on acute optogenetic stimulation as the valenced or reinforcing event and therefore do not have direct counterparts in constitutive NPF knockdown. We therefore focused the knockdown analysis on feeding and mating, which can be measured without imposed neural stimulation. Because NPF signaling has established roles in energy balance, and because cell-type-specific activity manipulations produced distinct changes in food intake, we also measured triglyceride and glycogen stores to determine whether functional specialization among NPF cell types extends beyond behavior to metabolic physiology.

Knocking down NPF in DM neurons reduced food intake, copulation rate, and glycogen levels (Figure 3A, B, D). Knockdown in L1 neurons reduced food intake, triglyceride, and glycogen (Figure 3A, C, D), whereas knockdown in P1 neurons selectively increased food intake (Figure 3A). Knockdown in P2 neurons selectively reduced glycogen (Figure 3D). The RNAi transgene reduced, but did not eliminate, NPF immunoreactivity in each cell type (Figure S2C); these phenotypes therefore identify functions that are sensitive to partial NPF reduction.

**Figure 3.**
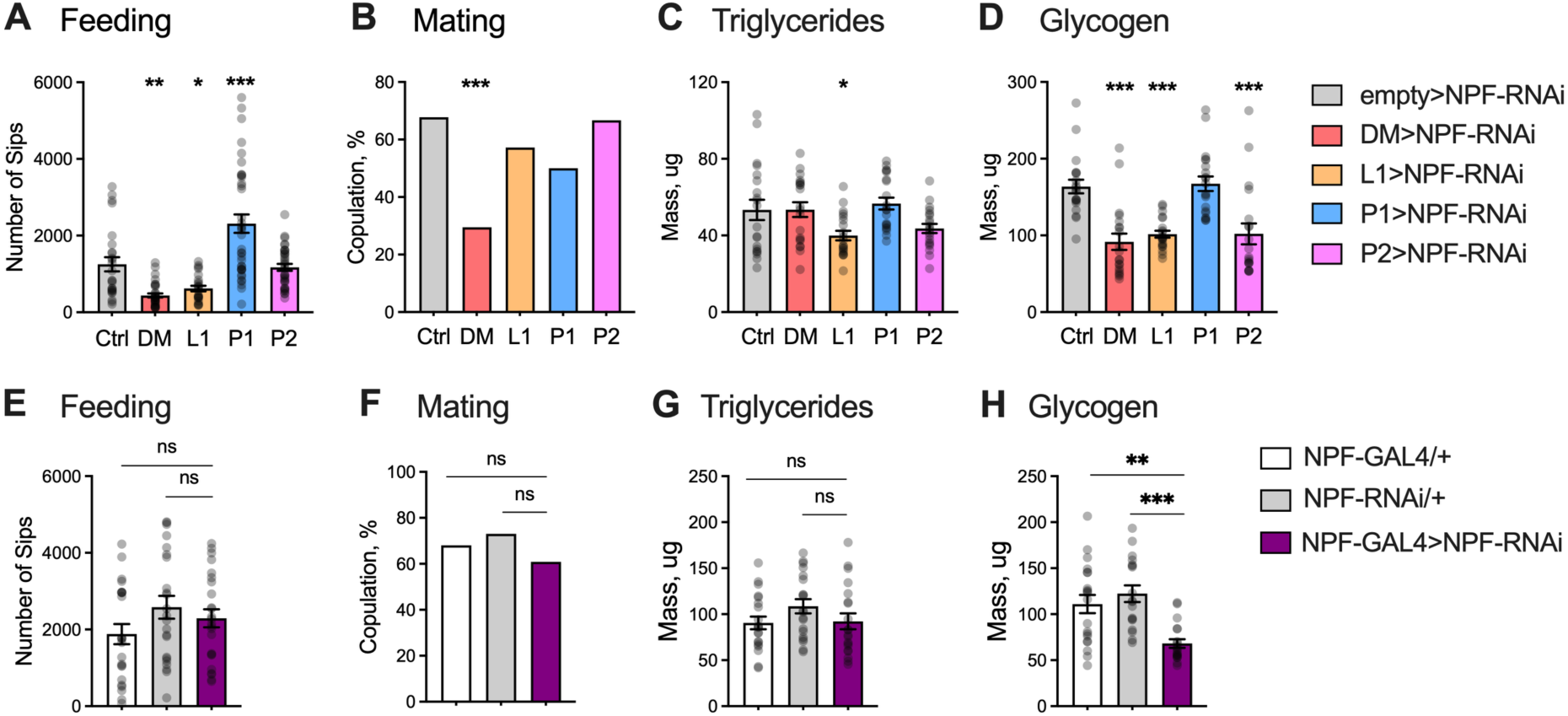
Cell-type-specific NPF knockdown reveals functions not apparent after global NPF loss. (A-D) Cell-type-specific knockdown of NPF produces distinct behavioral and metabolic phenotypes. NPF was knocked down in each subtype (DM, L1, P1, P2) and compared with the *empty>npf-RNAi* control (Ctrl). Knockdown in DM or L1 decreased food consumption, whereas knockdown in P1 increased it (A, feeding); knockdown in DM neurons reduced female copulation percentage (B, mating); knockdown in L1 selectively reduced triglyceride levels (C); and knockdown in DM, L1, and P2 reduced glycogen levels (D). (E-H) Knockdown of NPF in all NPF-expressing cells (*NPF-GAL4>NPF-RNAi*) did not affect feeding (E), mating (F), or triglyceride levels (G), but reduced glycogen levels (H) relative to both parental controls. N = 20-23 (feeding), n = 29-36 (mating), 20 (triglyceride and glycogen). Data are mean ± SEM. Statistics: Fisher’s exact test (mating); one-way ANOVA Dunnett’s multiple comparisons test (cell-type specific knockdown, feeding, triglyceride and glycogen) or Tukey’s multiple comparisons test (global knockdown, feeding, triglyceride and glycogen). \**p* < 0.05, \*\**p* < 0.01, \*\*\**p* < 0.001, ns. *p* > 0.05.

Thus, reducing NPF in different cell types produced distinct, partially overlapping behavioral and metabolic phenotypes.

We next asked whether these cell-type-specific phenotypes remain evident after population-level NPF loss. Pan-NPF knockdown using NPF-GAL4, which labels central neurons and gut enteroendocrine cells, did not detectably alter feeding, mating, or triglycerides but reduced glycogen relative to both parental controls (Figure 3E-H).

Glycogen was reduced after NPF knockdown in DM, L1, and P2 individually and after pan-NPF knockdown, indicating that several cell types make directionally aligned contributions to this metabolic readout. By contrast, the feeding, mating, and triglyceride phenotypes revealed by cell-type-specific knockdown were absent after population-level knockdown. Feeding provides a direct example of how opposing cell-type-specific effects can be obscured: NPF knockdown in DM or L1 decreased intake, whereas knockdown in P1 increased it.

Overall, population-level NPF loss produced a substantially narrower phenotype than cell-type-specific knockdown. The results are consistent with opposing, compensatory, or nonlinear interactions among cellular sources and demonstrate that population-level peptide loss does not provide a complete inventory of cell-type-specific NPF functions.

### NPF peptide contributes conditionally to phenotypes induced by pan-NPF activation

Activity manipulation and peptide knockdown therefore provide complementary but non-equivalent views of the NPF system (Figures 2 and 3). Because the two perturbations addressed different questions, the assay sets were not fully symmetric. We examined positional preference and appetitive conditioning only with activity manipulation, where optogenetic stimulation served as the valenced (i.e., reinforcing) event. For peptide knockdown, we focused on feeding and mating, which can be measured without imposed neural stimulation, and additionally measured triglyceride and glycogen stores because NPF has established roles in energy balance and cell-type-specific activity manipulation produced distinct feeding phenotypes.

Comparing these complementary maps revealed an asymmetry in how cell-type-specific effects were represented at the population level (Figure 4A). Population-level activity manipulation reproduced the direction of cell-type-specific effects in positional preference, learning, and feeding, but not mating, whereas population-level NPF knockdown retained only the glycogen phenotype (Figure 4A). Notably, glycogen was the only readout for which knockdown in multiple cell types shifted the measure in the same direction; the cell-type-specific feeding, mating, and triglyceride phenotypes were not evident after pan-NPF knockdown. This contrast indicates that the relationship between cell-type-specific and population-level phenotypes depends on whether the aggregate neuronal output or NPF peptide alone is perturbed. We therefore asked directly how much NPF contributes to the robust behavioral effects of pan-NPF activation.

**Figure 4.**
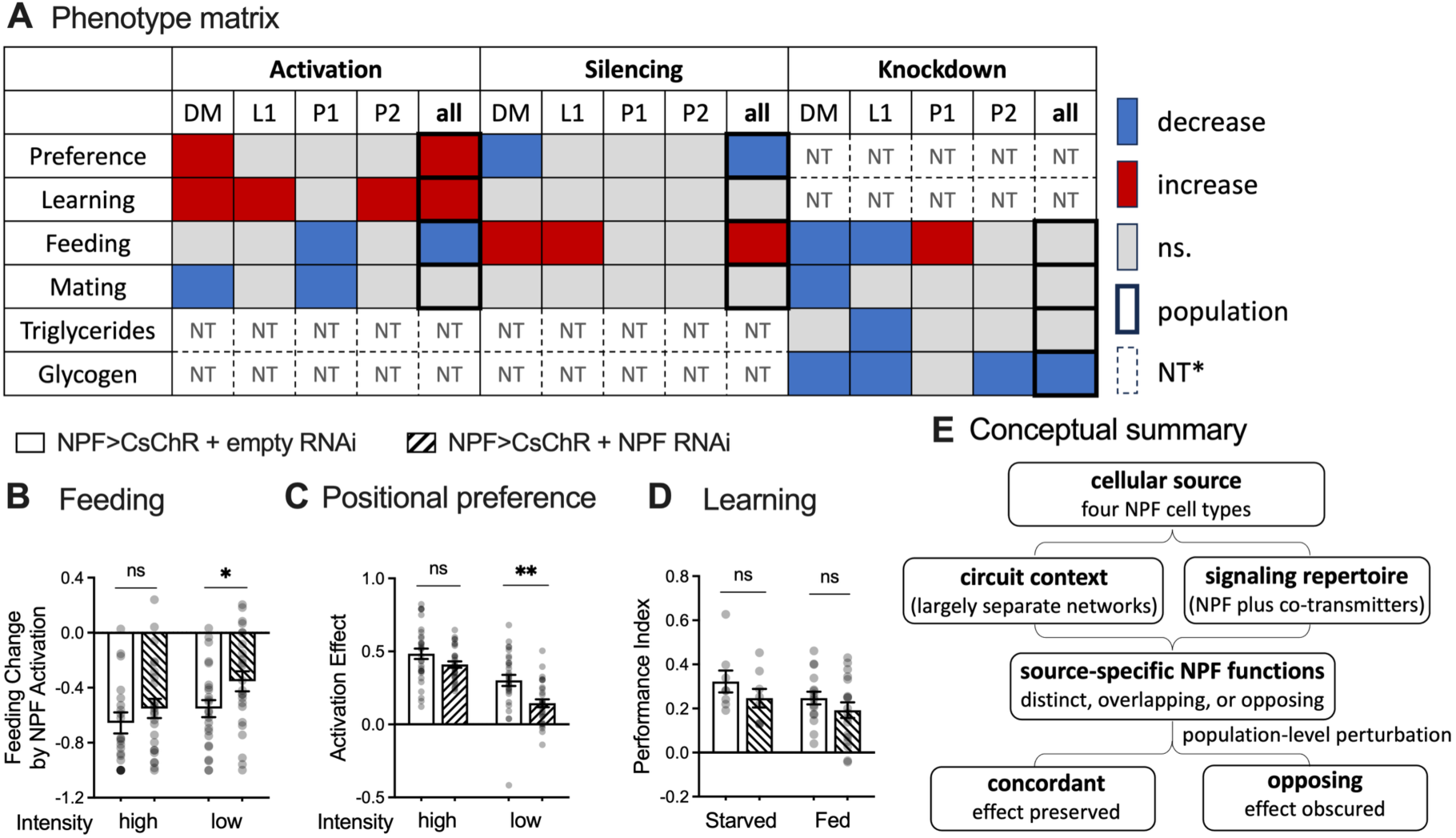
NPF peptide contributes partially to the phenotypes driven by NPF-neuron activity. (A) Summary matrix comparing phenotype direction after cell-type-specific or global neuroactivity manipulation and NPF knockdown. Across the assays tested, global activity manipulation recapitulated the direction of most cell-type-specific activity phenotypes, whereas global NPF loss generally failed to reveal phenotypes observed after cell-type-specific knockdown. Mating and glycogen were the respective exceptions. Gray indicates no detectable effect, dashed boxes indicate conditions not tested, and outlined boxes indicate global manipulation. * Gray cells marked “NT” indicate assays that were not performed, rather than an absence of phenotype. Positional preference and olfactory conditioning were examined only in the activity-manipulation series because neural stimulation served as the valenced or reinforcing event. Triglyceride and glycogen were examined only in the knockdown series to assess sustained peptide-dependent effects on energy storage and were not included in the acute activity-manipulation series. (B) During global NPF-neuron activation, simultaneous NPF knockdown did not detectably alter feeding suppression at the higher stimulation intensity (12 mW/cm²) but attenuated suppression at the lower intensity (6 mW/cm²). (C) During global NPF-neuron activation, simultaneous NPF knockdown did not significantly alter positional preference at the higher stimulation intensity (1 mW/cm²) but attenuated preference at the lower intensity (0.6 mW/cm²). (D) Simultaneous NPF knockdown did not detectably alter appetitive memory generated by pairing global NPF-neuron activation (1 mW/cm²) with an olfactory cue in either fed or starved flies. (E) cellular source and circuit context organize NPF function within a single-receptor system. Four NPF cell types constitute distinct cellular sources that differ in circuit context (largely separate input–output networks; Figure 1) and signaling repertoire (NPF plus other activity-dependent signals; Figures 2 and 4). These generate source-specific NPF functions that are distinct, overlapping, or opposing (Figures 2 and 3). Population-level perturbation collapses them into a composite readout in which concordant contributions remain detectable, whereas opposing or source-restricted contributions do not. “Obscured” denotes an undetectable effect, not a demonstrated cancellation.

To address this question, we combined pan-NPF activation with *npf* RNAi and tested the three assays in which global activation produced a phenotype: feeding, positional preference, and olfactory learning. For feeding, results above showed that global activation suppressed fly’s food intake (Figure 2I). We quantified the activation-induced change relative to the no-LED baseline as (Sip_LED_ − Sip_NoLED_)/Sip_NoLED_, where −1 denotes complete suppression and 0 denotes no change. NPF knockdown did not detectably alter activation-induced feeding suppression at the higher stimulation intensity but attenuated it at the lower intensity (Figure 4B). Similarly, NPF knockdown attenuated activation-induced positional preference at the lower, but not the higher, stimulation intensity (Figure 4C). In olfactory conditioning, however, NPF knockdown did not detectably reduce appetitive memory generated by high-intensity pan-NPF activation in either fed or starved flies (Figure 4D). We were unable to test lower-intensity reinforcement because weaker stimulation would shift memory toward the assay floor and reduce sensitivity to genotype-dependent differences.

Together, these experiments show that NPF peptide makes a conditional rather than uniform contribution to the behavioral consequences of activating the NPF neuron population. Its contribution was detectable for feeding suppression and positional preference under weaker stimulation intensities, whereas these phenotypes at higher intensity—and appetitive memory under the conditions tested—remained intact after NPF knockdown. The lack of an effect at high intensity does not establish that NPF is dispensable, because strong activation may recruit sufficient NPF-independent output to saturate the behavioral response and thereby mask a peptidergic contribution. Thus, the principal conclusion is that pan-NPF activation cannot be interpreted simply as NPF release: NPF mediates a context- and intensity-dependent component of the response, while additional activity-dependent signals are likely to contribute to the remainder.

## Discussion

A neuropeptide system that signals through only one receptor can be nevertheless partitioned into functionally distinct and sometimes opposing cellular functions. This is the principal finding of our study (Figure 4E). By resolving four sexually monomorphic adult NPF cell types and perturbing each at the levels of neuronal activity and peptide expression, we show that DM, L1, P1, and P2 occupy largely separate networks and make separable contributions to reward-related behavior, feeding, reproduction, and metabolism. Reducing NPF within individual cell types uncovered peptide requirements that were mostly absent after population-level perturbation, whereas manipulating neuronal activity revealed functions not explained by NPF alone. Thus, the functional breadth of the NPF system does not arise from a homogeneous population broadcasting a common message. It emerges from source-defined modules that share NPF and NPFR but differ in connectivity, behavioral role, and signaling repertoire.

### Cellular source creates specificity within a single-receptor peptide system

Current frameworks of peptidergic neuromodulation propose that peptide function must be resolved at the level of the producing cells and circuits ^2,3^. Our results provide a cell-type-resolved test of this principle within one peptide system in which receptor multiplicity cannot explain functional diversification.

The four NPF cell types had largely distinct first-order partners, with only a single presynaptic partner shared between DM and L1, and showed only modest second-order similarity, placing them in largely separate input-output networks (Figure 1). Their functions were likewise distinct but partially overlapping. DM and L1 were the most similar pair in the connectome, yet their functional profiles remained separable: both contributed to NPF-dependent control of feeding and glycogen levels, whereas DM additionally affected copulation success and L1 affected triglyceride levels (Figure 3). Circuit position therefore provides a structural basis for source-dependent functional specialization without implying complete independence: NPF cell types may still be functionally linked through shared neuropils and higher-order partners, and NPF released from one source may reach extrasynaptic targets beyond its direct synaptic network.

### Population-level perturbation can obscure cell-type-specific functions

Population-level activity manipulation preserved the direction of cell-type-specific effects in positional preference, appetitive learning, and feeding, but not in mating success (Figure 2). Even within feeding, different cell types appeared to dominate the population-level outcome in the two directions: activating all NPF neurons decreased intake, matching P1 activation, whereas silencing all NPF neurons increased it, matching DM or L1 silencing. Strong or directionally compatible source-specific effects can therefore remain visible at the population level, although the mating result shows that this is not guaranteed.

Population-level peptide perturbation, however, produced a substantially narrower profile (Figure 3). NPF knockdown in DM or L1 decreased feeding, whereas knockdown in P1 increased it; pan-NPF RNAi did not produce a detectable feeding phenotype. By contrast, reducing NPF in DM, L1, or P2 shifted glycogen levels in the same direction, and reduced glycogen remained detectable after population-level disruption. These findings show that global peptide perturbation can obscure cell-type-specific functions, particularly when different cellular sources make opposing contributions, whereas directionally aligned effects are more likely to remain detectable.

This pattern should not be necessarily interpreted as simple cancellation. Perturbation efficacy (RNAi did not eliminate the peptide), developmental compensation, nonlinear interactions, and peripheral NPF sources may also shape the phenotype. Thus, a population-level peptide phenotype is a composite readout rather than an inventory of cell-type-specific function. This finding provides an experimental example of the “conglomerate phenotype” problem proposed for broadly distributed neuropeptide systems ^2^.

### Neuronal activity is not equivalent to peptide action

Activity manipulations test the aggregate output of NPF neurons, whereas *npf*-RNAi tests the specific contribution of NPF while leaving the rest of the neuronal output intact. Agreement between the two manipulations supports a peptide contribution; divergence may reflect co-released signals, interactions among outputs, or differences in perturbation efficacy and timescale.

In DM and L1, neuronal silencing increased feeding whereas NPF reduction decreased it (Figures 2E and 3A). These divergent effects show that neuronal silencing and peptide depletion are not interchangeable. They are consistent with DM/L1-derived NPF promoting intake while the aggregate activity-dependent output of these neurons suppresses it. In P1, by contrast, activation decreased feeding whereas NPF reduction increased it, supporting a contribution of P1-derived NPF to feeding suppression.

Combining pan-NPF activation with NPF knockdown directly tested whether NPF contributes to phenotypes produced by activating the entire NPF population (Figure 4B–D). If the activation phenotype depends in part on NPF, reducing the peptide should attenuate the response. Reducing NPF attenuated feeding suppression and positional preference at the lower stimulation intensity, supporting a peptide contribution to both responses. The same knockdown did not detectably alter either phenotype at higher intensity and did not reduce appetitive memory at the high intensity used for that assay.

These results do not indicate that NPF acts only at low stimulation intensity. Rather, stimulation intensity determines whether its contribution is experimentally detectable. Under weaker activation, partial loss of NPF is behaviorally consequential. Under stronger activation, residual peptide (Figure S2C) may be released in sufficient amounts, or stronger activation may recruit enough co-released signals to preserve the response despite reduced NPF. These mechanisms are not mutually exclusive and cannot be distinguished here, but both explain why high-intensity activation can conceal the peptide contribution that is detectable at lower intensity.

Together, these experiments establish that NPF is a genuine contributor to the behavioral output: reducing the peptide attenuates phenotypes produced by activating the neurons that release it. They also show that the peptide is not account for the neuron’s complete output: under strong activation, the same reduction had no detectable effect. Residual peptide, co-released signals, or both may preserve the response. Activating an NPF neuron is therefore not equivalent to releasing NPF, and how far the two diverge depends on how strongly the neuron is stimulated.

### Cell-type specialization reframes an apparent NPF reward paradox

Cell-type specialization provides a framework for reconciling an apparent tension in the NPF literature. In food consumption, the NPF system is engaged when food reward is lacking: NPF promotes food seeking and permits hunger-dependent appetitive memory expression ^11,12,15^. In sexual experience, the NPF system is engaged when reward has been obtained: copulation increases brain NPF, whereas sexual deprivation reduces NPF-system activity and increases ethanol seeking; activating NPF neurons is itself reinforcing and reduces subsequent ethanol seeking ^18,19^. Thus, the NPF system has been linked both to pursuing an absent reward and to the state following reward attainment. These studies differ in sex, developmental stage, cell population, and assay, but together they resist a single unitary interpretation of NPF as either a deprivation or attainment signal.

The reinforcement data provide initial evidence that distinct NPF cell types can recruit different reward-related outputs. DM and P2 activation recruited PAM dopaminergic compartments and supported appetitive memory, whereas L1 supported appetitive memory without detectable recruitment of the PAM compartments examined or immediate positional preference (Figure 2), a pattern consistent with unsampled PAM pathways, subthreshold recruitment, or a distinct reinforcement route. The previously described NPF-PPL1 pathway provides one precedent for NPF regulation of an alternative dopaminergic pathway ^27^, although its relationship to L1 remains unresolved. Thus, NPF cell types can access reinforcement through more than one circuit architecture.

Resolving the apparent reward paradox will require measuring natural activity in defined NPF cell types during hunger, feeding, sexual deprivation, and copulation, and determining whether NPF and co-released signals act in the same or opposite directions in each pathway.

### A single-receptor architecture with a mammalian counterpart

Together, these findings establish cellular source as a determinant of NPF function at the levels of circuit position, population phenotype, and the relationship between neuronal activity and peptide action. This principle may extend to other peptide systems in which diverse sources signal through a shared receptor.

Mammalian NPY provides a different context. NPY is produced by hypothalamic AgRP neurons, cortical and hippocampal interneurons, brainstem catecholaminergic populations, and other cell types embedded in distinct circuits ^5,36–41^; its actions are further diversified by multiple receptor subtypes ^8^. Our study provides a tractable experimental demonstration that cellular diversity can map onto distinct circuit positions and separable behavioral and physiological functions within a single NPY-family system.

*Drosophila* NPF isolates the contribution of cellular source because it signals through a single known receptor, NPFR ^10^; known receptor-subtype identity therefore cannot partition its functions. The mammalian nociceptin/orphanin FQ (N/OFQ) system has a similar architecture as NPF: N/OFQ acts through the broadly distributed NOP/OPRL1 receptor, with no established receptor subtypes ^42^, while prepronociceptin precursor (Pnoc) is expressed in anatomically distinct neuronal populations with sharply different behavioral roles.

Three Pnoc populations illustrate this functional diversity. Central-amygdala Pnoc neurons are required for palatable food consumption and support reward ^43^; paranigral VTA Pnoc neurons constrain reward seeking through local NOP signaling ^44^; and extended-amygdala Pnoc neurons encode salient stimuli and promote defensive behavior ^45^. Thus, one peptide acting through one receptor can support functions that receptor identity alone cannot distinguish.

Furthermore, many studies of Pnoc-expressing neurons manipulate the cells’ aggregate output, which can include co-released classical transmitters and other peptides, leaving unresolved whether neuronal function can be equated with N/OFQ function. Our matched activity- and peptide-level perturbations show that neuronal function and peptide function can diverge (Figures 2-4). Thus, functional specificity in single-receptor peptide systems may reflect both source-specific circuit organization and the distinct signaling repertoire of each peptide-producing cell type.

### Limitations and future directions

The study examines four sexually monomorphic central cell types, not the entire NPF system. Synaptic connectomics cannot identify the complete set of extrasynaptic peptide targets. Characterizing sexually dimorphic and peripheral cell types, validating co-transmitters, mapping NPFR-expressing cell types, and combining cell-type-specific activation with adult-restricted peptide knockdown will be essential next steps.

Some of these unresolved questions are now experimentally tractable because of the cell-type-specific drivers developed here. Stable drivers for DM, L1, P1, and P2, validated anatomically and linked to their connectomic identities, permit these populations to be monitored and manipulated independently. Combined with temporal control and peptide-specific perturbations, these reagents provide a foundation for recording endogenous activity, identifying downstream NPFR-expressing targets, and distinguishing peptide-dependent from co-transmitter-dependent functions across physiological and behavioral states.

Together, these findings establish cellular source as an organizing principle of NPF signaling: one peptide acting through one receptor can produce distinct, and sometimes opposing, behavioral effects depending on the neurons from which it is released. By separating peptide action from the aggregate output of these neurons, our study further shows that peptidergic function is shaped jointly by source-specific circuit organization and the broader signaling repertoire of each cell type. Thus, understanding a peptidergic system requires defining not only the peptide and its receptor, but also which cells release it, which circuits they engage, and what other signals those cells convey.

## Materials and Methods

### Key Resources Table

| Reagent Type<br>(Species or Resource) | Designation | Source | Identifier | Assay |
| --- | --- | --- | --- | --- |
| Genetic Reagent<br>(*D. melanogaster*) | UAS-NPF-RNAi (I) | Shao Lab | VDRC:v108772 | All |
| Genetic Reagent<br>(*D. melanogaster*) | empty-split-GAL4 | Shao Lab | BDSC:79603 | All |
| Genetic Reagent<br>(*D. melanogaster*) | DM-spGAL4/TM6B | This Study | N/A | All |
| Genetic Reagent<br>(*D. melanogaster*) | L1-spGAL4 | Feng et al., 2021<sup>27</sup> |  | All |
| Genetic Reagent<br>(*D. melanogaster*) | P1-spGAL4 | This Study | N/A | All |
| Genetic Reagent<br>(*D. melanogaster*) | P2-spGAL4 | Shao et al., 2017<sup>19</sup> | N/A | All |
| Genetic Reagent<br>(*D. melanogaster*) | UAS-A1ACR-EYFP<br>(attP18) | Bushey et al., 2025<sup>30</sup> | N/A | Optogenetic Experiments |
| Genetic Reagent<br>(*D. melanogaster*) | UAS-CsChrimson-mVenus<br>(attP18) | Klapoetke et al., 2014<sup>29</sup> | BDSC: 55134 | Optogenetic Experiments |
| Genetic Reagent<br>(*D. melanogaster*) | UAS-Chrimson88-tdTomato (attP18) | Strother et al., 2017<sup>46</sup> | N/A | Calcium Imaging |
| Genetic Reagent<br>(*D. melanogaster*) | LexAop-CsChrimson88-tdTomato (attP18) | Y. Shuai | N/A | Calcium Imaging |
| Genetic Reagent<br>(*D. melanogaster*) | R58E02-LexA (attP40) | Pfeiffer et al., 2010<sup>47</sup> | BDSC: 52740 | Calcium Imaging |
| Genetic Reagent<br>(*D. melanogaster*) | LexAop-jGCaMP7s<br>(Su(Hw)attP8) | A. Wong | N/A | Calcium Imaging |
| Genetic Reagent<br>(*D. melanogaster*) | NPF-GAL4 | Shao Lab (Wu et al., 2003<sup>11</sup>) | N/A | All |
| Genetic Reagent<br>(*D. melanogaster*) | *w<sup>1118</sup>* | Shao Lab | N/A | All |
| Genetic Reagent<br>(*D. melanogaster*) | Elav-GAL4 | Shao Lab |  | IHC |
| Antibody | Rabbit anti-GFP | Invitrogen | A-6455;<br>RRID:AB_221570 | IHC |
| Antibody | Mouse nc82 | Hybridoma Bank DSHB | NC82;<br>RRID:AB_2314866 | IHC |
| Antibody | Rabbit anti-NPF | Ray Biotech, Inc | RB-19-0001 | IHC |
| Antibody | Mouse anti-GFP | Millipore Sigma | G6539 | IHC |
| Antibody | Alexa 488 goat anti-rabbit | Invitrogen | A-11008 | IHC |
| Antibody | Alexa 488 goat anti-mouse | Invitrogen | A-11001 | IHC |
| Antibody | Alexa 568 goat-anti-mouse | Invitrogen | A-11031 | IHC |
| Antibody | Alexa 568 goat-anti-rabbit | Invitrogen | A-11011 | INC |
| Chemicals | Normal goat serum | Gibco | 16210-064 | IHC |
| Chemicals | Paraformaldehyde, 20% aqueous solution | Electron Microscopy Science | 15713 | IHC |
| Chemicals | Phosphate buffer saline | Apex BioResearch | 18-244 | IHC |
| Chemicals | Schneider's Drosophila medium | Gibco | 21720-024 | IHC |
| Chemicals | Agar Powder | Thermo Scientific | 9002-18-0 | flyPad/optoPad |
| Chemicals | All trans-retinal | Spectrim Chemical Mfg. Co. | R3041-1GM | Optogenetic Experiments |
| Chemicals | Amyloglucosidase | Sigma-Aldrich | 10115-1G-F | Metabolism |
| Chemicals | 0.1 M Citric Acid | Fisher | 5949-29-1 | Metabolism |
| Chemicals | D-Sucrose | Fisher BioReagents | BP220-212 | flyPad/optoPad |
| Chemicals | Glucose Standard Solution | Sigma-Aldrich | G6918 | Metabolism |
| Chemicals | Glycerol Standard Solution | Sigma-Aldrich | G7793 | Metabolism |
| Chemicals | Infinity Triglyceride Reagent | Fisher BioReagents | TR22421 | Metabolism |
| Chemicals | 5 M NaCl | Sigma-Aldrich | 7647-14-5 | Metabolism |
| Chemicals | Pointe Scientific Glucose Oxidase | Fisher BioReagents | 23-666-290 | Metabolism |
| Chemicals | Protease Inhibitor | Sigma-Aldrich | 04 693 124 001 | Metabolism |
| Chemicals | 0.1 M Sodium Citrate | Sigma-Aldrich | 6131043 | Metabolism |
| Chemicals | 1 M Tris-HCl | KD Medical | RGE-3340 | Metabolism |
| Chemicals | Triton-X | Sigma-Aldrich | 9002-93-1 | Metabolism |
| Chemicals | Yeast Flakes | Any Lab Supplier | N/A | Choice |
| *Drosophila* Medium | Nutri-fly Molasses Formulation Fly Medium | Genesee Scientific | 66-123<br>https://geneseesci.com | All |
| Equipment | flyPAD and optoPAD High Throughput Feeding Assay | Itskov et al. 2014 | https://flypad.rocks/ | flyPad/optoPad |
| Equipment | Microplate Reader | N/A | N/A | Metabolism |
| Equipment | Modular In-Vivo Multiphoton Microscopy System (MIMMS) v2.0 | Janelia Research Campus, HHMI | https://www.janelia.org/open-science/mimms-22-2024;<br>RRID:SCR_016511 | Calcium Imaging |
| Software, Algorithms | BioRender | BioRender | https://biorender.com | Data Analysis |
| Software, Algorithms | GraphPad Prism 10 Software | GraphPad Software | https://www.graphpad.com/features | Data Analysis |
| Software, Algorithms | MATLAB R2024a software | The Mathworks, Inc. | https://www.mathworks.com/ | Data Analysis |
| Software, Algorithms | Python Programming Language Software | Python Software Foundation | https://www.python.org/;<br>RRID:SCR_008394 | Mating; Connectome Analysis |

### Fly stocks and culture

*Drosophila melanogaster* stocks (listed in key resources table) were raised at 25°C and 50%-60% relative humidity on commercially prepared standard media (cornmeal/agar/molasses/yeast; Nutri-Fly MF) for anatomical and ‘Con-Ex’ feeding experiments. For optogenetic behavioral and *ex vivo* functional imaging experiments, flies were raised under constant darkness at 25°C and 50%-60% relative humidity on the same media supplemented with 0.2 mM all trans-retinal. Flies were collected 0-3 days post-eclosion on media containing 0.4 mM all trans-retinal. Flies for anatomical and behavioral assays were 4- to 7-days-old at the time of experiments; flies used for *ex vivo* functional connectivity calcium imaging were 2- to 5-days-old at the time of dissection. Mated female flies were used in this study, with the exception of copulation experiments, where unmated female flies were used.

### Connectome analysis

Connectivity of NPF subtype neurons was analyzed using the FlyWire/Codex whole-brain connectome of the FAFB (snapshot 783). Custom Python code, adapted from the analysis framework of Walker et al., 2025 ^48^, was used to extract each subtype’s direct (first-order) and second-order pre- and postsynaptic partners, applying minimum synapse-count thresholds of 5 and 10 synapses, respectively, to exclude weak connections. Partner overlap among subtypes was visualized with Venn diagrams and quantified using the Jaccard similarity index; the significance of pairwise partner overlap was assessed by hypergeometric tests with Benjamini-Hochberg false discovery rate correction, and differences between input- and output-partner sharing were evaluated using Fisher’s exact test and a paired Wilcoxon signed-rank test across subtype pairs. Additional analyses characterized each subtype’s neurotransmitter composition, neuropil innervation profile, downstream/upstream partner super-class composition, cosine similarity of synapse-weighted partner vectors, and input/output synapse polarity. All analysis code is available at https://github.com/lishashao/NPF-subtype-connectome-analysis.

### Behavioral assays

#### Positional preference

Positional preference assays were performed as described previously at 25°C, 50%-60% relative humidity in darkness^19^. Female flies (3-7d old) were loaded into rectangular chambers, 8 flies per chamber. After a 2-min baseline period, one side of the chamber (selected at random) was illuminated by 0.5 Hz 1-second pulsed 625-nm red LED at designated light intensity (indicated in figure legends) for 3 min. Following the illumination period, there was a 2-min recovery period. Experiments were recorded with infrared cameras (BlackFly USB3 camera, Teledyne vision solutions). The frequencies and intensities of LED stimulation were controlled using MATLAB (MathWorks, Inc.). Acquired video was analyzed with custom MATLAB scripts that detected the position of the flies, allowing for the calculation of a Preference Index as the average difference in the fractions of flies on illuminated versus unilluminated sides.

#### Associative learning

Associative learning assays were performed on female flies with optogenetic stimulation and were conducted in a circular arena that couples optogenetic stimulation with an odor delivery system, as described in Aso et al., 2016 ^28^. All tests were performed at 25°C, 50%-60% relative humidity. The olfactory cues used were 4-methylcyclohexanol (1:1000) and 3-octanol (1:1000). Optogenetic stimulation used a 0.5 Hz 625-nm LED light at designated intensity (indicated in figure legends). In optogenetic learning assay, female flies were trained as groups of 12-14 individuals using a single training session consisting of 1-min exposure to air, followed by a 1-min exposure to odor A (CS+) paired with neuronal activation/silencing, followed by a 1-min exposure to air, and finally a 1-min exposure to odor B. Memory was tested 1 min after training, during which the odors A and B were delivered to each pair of opposing quadrants. Experiments were recorded under IR illumination with a camera (Point Gray Research) located above the chamber and equipped with an IR long-pass filter. The subsequent video was analyzed with custom MATLAB scripts that detect the position of flies within the chamber. A Performance Index for odor A (paired with LED, CS+) is the average difference in the fractions of flies on the odor-A versus odor-B quadrants. A reciprocal group received odor B as CS+ (paired with LED) to counterbalance flies’ intrinsic odor bias. The final Performance Index is the mean of PI(A-CS+) and PI(B-CS+).

#### Mating

The courtship assay assessed copulation behavior between a virgin experimental female fly and a sexually mature CantonS (CS) male fly. Individual female and male flies (4-7d old) were loaded separately using a fly aspirator into a small circular chamber (1-cm in diameter). The flies were initially separated by a thin movable barrier to prevent premature interaction. At the start of the assay, the barrier was removed and behavior was recorded for 70 minutes. In optogenetic experiments, chambers were illuminated with 625-nm red LED light delivered at 1s intervals to stimulate channelrhodopsin-expressing neurons. Mating behavior was analyzed using Fly Flirt Python-based software, which detected copulation events, including copulation onset and duration. Flies that did not mate during the recording period were also identified by the software and included in data analysis.

#### flyPad/optoPad feeding

This assay was performed as described previously by using the flyPad behavioral monitoring system,25,26 at 25°C, 50% −60% relative humidity ^35^. Warmed, liquefied media were loaded into both probes in each FlyPAD chamber in darkness. Individual female flies (3-7 d old) were placed in each flyPAD chamber and were allowed to feed ad libitum for 50 min total. In the open-loop feeding and choice assay, the flyPAD chambers were illuminated with 625-nm red LEDs to optogenetically stimulate channelrhodopsin-expressing neurons. In the closed-loop feeding assay, the FlyPAD chambers were only illuminated with red LED when flies fed on one of the two probes in each arena. The probe associated with illumination was alternated for the next set of experiments to mitigate potential probe-specific bias. Media containing 7% yeast, 160 mM sucrose, and 0.7% agar was used for open-loop feeding assay. Media containing either 0.7% agar (neutral taste), 100 mM sucrose (sweet taste), or 0.5 mM quinine (bitter taste) were used for open-loop choice assays and closed-loop feeding assays. Analysis of feeding behavior used custom MATLAB scripts that extracted the total number of sips during the feeding period.

### Metabolites measurement

Metabolic activity was assessed by quantifying whole-fly triglyceride, glycogen, and free glucose levels using a protocol adapted from Murakami et al., 2016 ^49^. Two female flies (3-7d old) were euthanized and homogenized in 250 μL of lysis buffer supplemented with protease inhibitors at a 7X concentration. Samples were centrifuged at 13,000 RPM for 15 minutes at 4°C and the resulting supernatant (∼200 μL) was used for further metabolite assays. For each metabolic assay, 10 µL of the supernatant was loaded in triplicate into a 96-well plate. To quantify triglycerides, 90 μL of infinity triglyceride reagent was added to the sample-filled wells. Similarly, to quantify free glucose, 90 μL of glucose oxidase reagent was added to additional sample-filled wells on a separate 96-well plate. Reagents were stored at 4°C and incubated with the samples at 37°C for 5 minutes with shaking. To measure glycogen, 20 μL of amyloglucosidase was added to 20 μL of supernatant to hydrolyze the *α*-1,4 glycosidic bonds, followed by incubation at 37°C for 2 hours. After incubation, samples were loaded in triplicate into 96-well plate, and 90 μL of glucose oxidase reagent was added. Absorbance was measured at 500 nm for all assays. Metabolite concentrations were calculated via standard curves and then converted to usable mass values for data analysis.

### Immunostaining and confocal imaging

Immunohistochemical staining and imaging were performed as follow. Briefly, female fly brains and guts were dissected in cold Schneider’s Insect (S2) medium, fixed in 2% paraformaldehyde in S2 at Room Temperature (RT) under nutation for 55-65 min in covered Eppendorf tubes, washed 3-4 times with PBT0.5 (phosphate buffered saline, PBS, containing 0.5% Triton X-100) at RT, blocked with PAT3 (1xPBS, , 3% Normal Goat Serum, 0.5% Triton X-100) for 1 hour at RT, incubated with primary antibody (1:500 rabbit anti-GFP, 1:50 mouse anti-nc82 in PAT3) or (1:250 mouse anti GFP and rabbit anti-NPF Supplementary figure)at 4°C overnight, washed 3-4 times in PBT0.5, incubated with secondary antibody (1:500 Alexa 488 anti-rabbit, 1:500 Alexa 568 anti-mouse in PAT3) or (1:500 Alexa 488 anti-mouse, 1:500 Alexa 568 anti-rabbit for NPF staining)) at 4°C overnight, washed with PBT0.5 and PBS, and mounted in Vectashield. For control of NPF staining when the anti-NPF were omitted while both secondary present, the specific staining was absent.The brains were then imaged on a confocal microscope (Zeiss CellDiscoverer 7, using Zen Blue software, Zeiss LSM880, using Zen Black software).

### *Ex vivo* calcium imaging

All functional imaging was performed on 2- to 5-day-old female flies. Whole brains were dissected in a Sylgard silicone elastomer (Dow Inc.) coated dish, immersed in artificial fly hemolymph (103 mM NaCl, 3mM KCl, 2 mM CaCl2, 4 mM MgCl2·6 H2O, 26 mM NaHCO3, 1 mM NaH2PO4, 8 mM trehalose, 10 mM glucose, 5 mM TES, bubbled with 95% O2/5% CO2). Brains were placed on a poly-Lysine coated coverslip and imaged with a resonant-scanning 2-photon microscope (MIMMS v2.0, Janelia Research Campus), using a laser with a 920 nm excitation wavelength (Toptica FemtoFiber Ultra 920) and an excitation power of 19-33 mW (2.7-4.3 mW/mm2 intensity). Emitted fluorescence was detected with GaAsP photomultiplier tubes (Hamamatsu) controlled via a Janelia PMT Controller. Images were acquired with an Olympus 40x water-dipping, 0.8 numerical aperture objective at 512 pixels × 512 pixels resolution. Optogenetic stimulation of CsChrimson was provided by a 660-nm red light LED diode (ThorLabs M660L4), fed into a digital mirror device (Texas Instruments DLP Lightcrafter 4500) and projected through the objective. Photostimulation consisted of 4 x 1 s pulses, separated by a 24 s inter-pulse interval, with a stimulation power of 20-21 μW (2.8-3.0 μW/mm2 intensity). Volume stacks were continuously imaged at 2x magnification, and consisted of 45 slices, with an inter-slice interval of 2 μm, thus allowing all the PAM-DANs compartments to be imaged. Stack acquisition began 10s prior to the 1st LED pulse and stopped 24s after the termination of the final pulse. After optogenetic stimulation, brains were imaged for ROI definition of mushroom body compartment anatomy using the same imaging stack dimensions as above, using both 920 nm, and 1050 nm excitation (Toptica FemtoFiber Ultra 1050). Then power intensity for imaging at the 920- and 1050-nm excitation was 3.2-3.4 and 3.1-3.2 mW/mm2 respectively.

Regions of interest (ROI)s for each brain were drawn using custom MATLAB software and the 1050 nm anatomy images described above. The average pixel intensity for each ROI (*fluor_avg_*) was calculated, as was its average baseline intensity (*fluor_bseln_*), defined as the averaged intensity for the 5 volumes (approximately 1.5 s) acquired immediately prior to each optogenetic pulse onset. Average background intensity (*fluor_bckgr_*) for each volume was calculated from a ROI located in non-brain-tissue containing space. For each PAM compartment, ΔF/F was calculated using the formula:

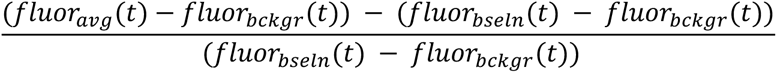

### Statistical analysis

Analyses were performed at the level of the experimental unit. For single-fly assays, the unit of analysis was the individual fly, such that each fly contributed one value (thus *N* = number of flies). For group assays (e.g., arena-based population tests), the unit of analysis was an independently loaded cohort of flies tested together in a single chamber/arena; all flies in that cohort were aggregated to yield one summary value (e.g., PI or total sips), and that value contributed *N* = 1. In these group assays, the number of flies per cohort (*n*) is reported descriptively in figure legends but was not used for statistical inference to avoid pseudo-replication. When cohorts were measured more than once under different conditions, within-cohort comparisons were analyzed with paired tests; otherwise, unpaired tests were used across independent cohorts.

For each dataset (and, where applicable, model residuals for >2-group designs), distributional normality was assessed with the Shapiro-Wilk test (α = 0.05). All hypothesis tests were two-tailed. When the normality criterion was met, two-group comparisons used paired or unpaired *t*-tests as appropriate to the design, and >2-group comparisons used one-way ANOVA followed by Tukey’s multiple-comparisons test. When the normality criterion was not met, the behavioral index was tested against 0 using a one-sample Wilcoxon signed-rank test; two-genotype comparisons used the Mann-Whitney U test; and >2-genotype comparisons used the Kruskal-Wallis test with Dunn’s post hoc multiple-comparisons procedure.

The exact test used, sample size (*n*), and two-tailed *p*-values are provided in the figure legends. Analyses were performed in GraphPad Prism.

## Acknowledgements

We thank Yoonwoo Park and Sabina Knox for their contributions to the early stage of this project. We thank Drs. Ulrike Heberlein, Amber Krauchunas, Jeremy Bird, and Jon-Michael Knapp for discussion and feedback. We thank We thank Dr. Meet Zandawala for valuable feedback on cell-type identification and technical assistance. We thank Drs. Ann-Shyn Chiang and Bloomington *Drosophila* Stock Center (BDSC) for providing fly strains. We thank Dr. Anita Devineni for sharing the code for connectome analysis and Mr. Sina Mirzaei for initial analysis. We acknowledge the Princeton FlyWire team and members of the Murthy and Seung labs for development and maintenance of FlyWire (supported by BRAIN Initiative grant MH117815 to Murthy and Seung). This work was supported by UD-GUR, UD-UDRF, Delaware CTR program with a grant from the National Institute of General Medical Sciences (NIGMS, U54 GM104941), Delaware INBRE program with a grant from the NIGMS (P20 GM103446), and Maximizing Investigators’ Research Award to L.S. (NIGMS R35GM147504).

## Data Availability

Requests for further information and resources should be directed to and will be fulfilled by the lead contact, Lisha Shao.

## Materials availability

Fly lines generated in this study are available upon request from the lead contact.

## Data and code availability

The code used for connectome analysis is described in Materials and Methods, and available at github.

Raw data for imaging and behavioral experiments are available upon request from the lead contact.

Any additional information required to reanalyze the data reported in this paper is available from the lead contact upon request.

