## Supplemental Figures and Tables for "Cellular source and circuit context organize functional specificity in the *Drosophila* NPF system"

### Supplementary Information

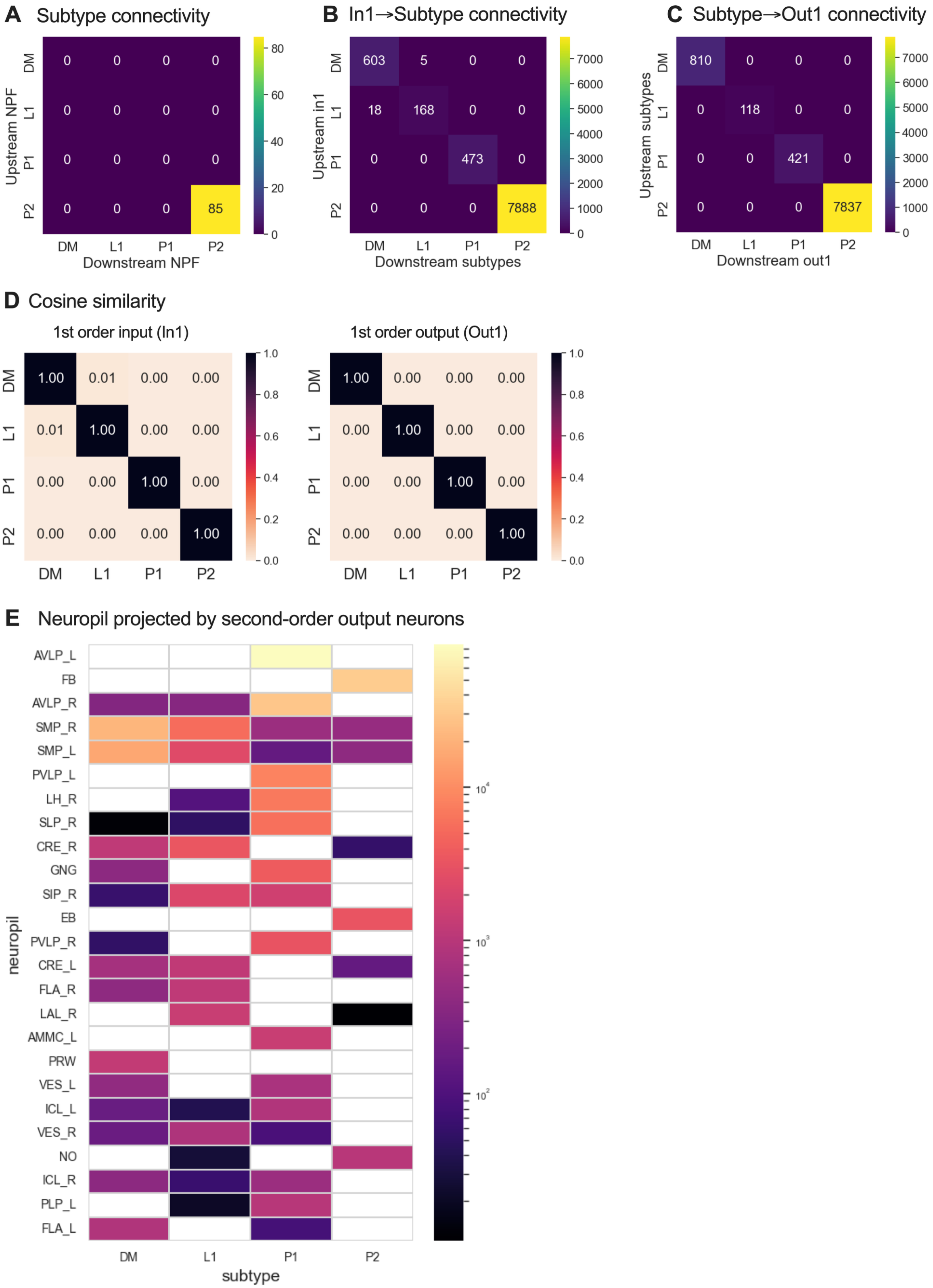

**Figure S1. NPF cell types and their synaptic partner populations are largely segregated in the connectome.**

(A) Synaptic connectivity among DM, L1, P1, and P2 neurons. No inter-type synapses were detected; P2 neurons formed within-type connections. Color indicates total synapse number.

(B) Connectivity from each first-order input population (In1) to the four NPF cell types. Each input population contacted its corresponding NPF cell type but not the other NPF cell types. Color indicates total synapse number.

(C) Connectivity from each NPF cell type to the four first-order output populations (Out1). Each NPF cell type contacted its corresponding output population but not the output populations assigned to the other NPF cell types. Color indicates total synapse number.

(D) Pairwise cosine similarity of synapse-weighted first-order input (In1; left) and output (Out1; right) vectors. Values range from 0 (no similarity) to 1 (identical vectors); diagonal values are 1 by definition.

(E) Neuropils innervated by second-order output neurons (Out2) associated with each NPF cell type. Color denotes projected connectivity on the logarithmic scale shown. Hemisphere suffixes are omitted from the abbreviation definitions. AMMC, antennal mechanosensory and motor center; AVLP, anterior ventrolateral protocerebrum; CRE, crepine; EB, ellipsoid body; FB, fan-shaped body; FLA, flange; GNG, gnathal ganglia; ICL, inferior clamp; LH, lateral horn; MB_CA, mushroom body calyx; NO, noduli; PLP, posterior lateral protocerebrum; PRW, prow; PVLP, posterior ventrolateral protocerebrum; SAD, saddle; SIP, superior intermediate protocerebrum; SLP, superior lateral protocerebrum; SMP, superior medial protocerebrum; VES, vest; WED, wedge. See also Figure 1.

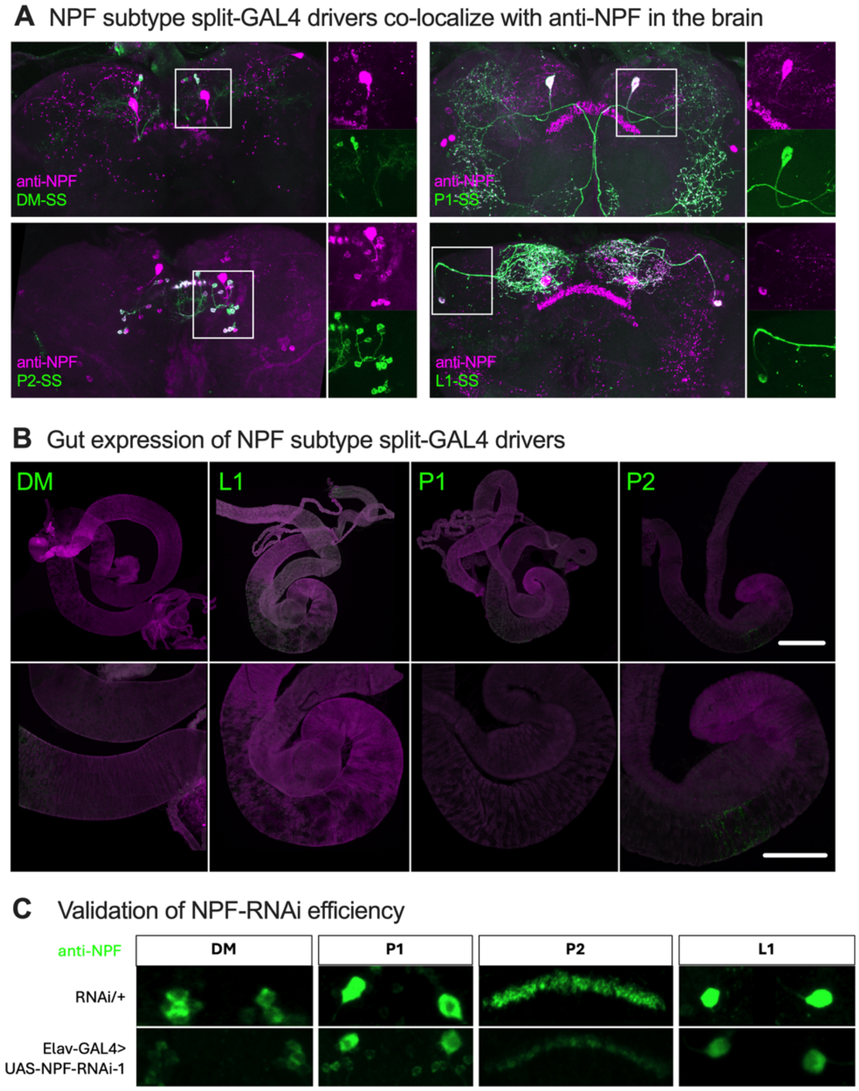


**Figure S2. Anatomical specificity of NPF cell-type drivers and validation of npf RNAi.**

(A) Co-localization of the DM, P1, P2, and L1 split-GAL4 driver patterns with endogenous NPF immunoreactivity in the adult brain. Driver-labeled neurons are shown in green and anti-NPF staining in magenta. Boxed regions are shown at higher magnification to the right of each image.

(B) Expression of the four NPF cell-type drivers in the adult gut. Representative whole-gut images are shown above, with higher-magnification views below. DM, L1, and P1 drivers showed little or no detectable peripheral expression under the conditions examined; the P2 driver showed weak labeling in a restricted gut region. Scale bars are 200uM (upper panels) and 100 uM (lower panels).

(C) Validation of the UAS-npf-RNAi-1 transgene by anti-NPF immunostaining. Representative DM, P1, P2, and L1 neurons are shown in UAS-npf-RNAi-1/+ controls (upper panel) and after panneuronal RNAi expression with elav-GAL4 (lower panel). RNAi reduced, but did not eliminate, NPF immunoreactivity across the four cell types.

**Table S1. FlyWire root IDs of NPF cell types included in the connectome analysis.**

Root IDs are provided for each reconstructed neuron assigned to the DM, L1, P1, or P2 cell type (in the right hemisphere) in the FAFB/FlyWire dataset used for the analyses in Figures 1 and S1. Each row represents one reconstructed neuron.

| NPF cell type | Root_ID |
|---|---|
| DM | 720575940622905383 |
| DM | 720575940624431460 |
| DM | 720575940625408133 |
| DM | 720575940632998114 |
| DM | 720575940631667281 |
| L1-l | 720575940627010234 |
| P1 | 720575940635549620 |
| P2 | 720575940606731529 |
| P2 | 720575940611117380 |
| P2 | 720575940612973096 |
| P2 | 720575940620932813 |
| P2 | 720575940622136129 |
| P2 | 720575940622899453 |
| P2 | 720575940625721331 |
| P2 | 720575940631483167 |
| P2 | 720575940631725087 |
| P2 | 720575940632718944 |
| P2 | 720575940633412971 |
| P2 | 720575940633744747 |
| P2 | 720575940634046399 |
| P2 | 720575940636914287 |
| P2 | 720575940641215104 |
